# Flow-Induced Vascular Remodeling On-Chip: Implications for Anti-VEGF Therapy

**DOI:** 10.1101/2024.11.29.625475

**Authors:** Fatemeh Mirzapour-Shafiyi, Elias Huber, Leonie Karr, Johnny Tong, Andreas R. Bausch, Friedrich C. Simmel, Karen Alim

## Abstract

Impaired vascular remodeling is linked to tumor progression, impacting the efficacy of anti-vascular therapies. However, the interplay between blood flow and vascular endothelial growth factor (VEGF) in driving vascular remodeling remains unclear. We optimized a human vasculature-on-chip system for long-term perfusion to quantify vascular remodeling under physiologically low and tumor-mimicking high VEGF conditions. Live imaging shows that at low VEGF levels, flow-conditioned vascular networks remodeled like animal models, enhancing flow velocities. Conversely, static networks lacking flow exhibited continuous growth, loss of network hierarchy, and decreased flow velocities. At tumor-mimicking high VEGF levels, flow-conditioned vessels showed aberrant overgrowth, which was blocked by the anti-VEGF tumor drug bevacizumab, restoring flow-induced remodeling. Without flow, however, high VEGF slowed vessel growth compared to low VEGF, while anti-VEGF treatment restored continuous growth. These findings provide valuable insights into optimizing anti-angiogenic therapies by exploiting VEGF modulation and flow conditions for vascular remodeling and improved tumor treatment.

## Introduction

Vascular remodeling is essential for maintaining vascular health and plays a key role in several diseases [1]. In tumor growth, vascular remodeling is often deficient, resulting in disorganized and dysfunctional vessels [2]. In contrast, diseases such as diabetes and aging are associated with excessive vascular remodeling, resulting in microvascular rarefaction, characterized by the loss or reduction of small blood vessels [3]. Vascular remodeling transforms the primitive vascular plexus - first formed from individual vascular precursor cells through 'vasculogenesis' and later expanded through 'angiogenesis', into a hierarchically organized, “normalized” network [1]. This transformation is critical for proper vascular function, as a normalized network is essential for efficient blood flow, nutrient delivery, and waste removal across tissues. Without effective remodeling, vascular networks cannot meet the demands of complex tissue environments, compromising tissue function and health [1]. Despite its critical importance, the precise mechanisms driving the interplay of factors that shape vascular remodeling, particularly under different pathological and physiological conditions, remain poorly understood.

Both the initial formation of new blood vessels from vascular precursor cells (vasculogenesis) and the sprouting of new vessels from existing vessels (angiogenesis) are driven by vascular endothelial growth factors (VEGFs), which promote endothelial cell proliferation, migration, and survival [4]. Among these, VEGF-A - particularly its most abundant and biologically active isoform, VEGF-A^164^ - plays a central role in regulating blood vessel formation in health and disease [5]. VEGF is released by tissue undergoing hypoxia and stimulates the formation of new blood vessels by vascular endothelial cells [4]. In the early stages of vessel development, elevated levels of VEGF stimulate rapid angiogenesis, followed by a maturation phase in which VEGF is downregulated [6]. While proper regulation of VEGF signaling is critical for the formation of functional blood vessels [6], its dysregulation, such as in the tumor environment, leads to abnormal vascularization [4], [7]. However, how VEGF signaling interacts with other factors, such as mechanical cues from blood flow, to control vascular remodeling lacks understanding, limiting our ability to effectively target these processes in disease contexts.

In addition to a regulated balance of VEGF signaling, blood flow is also critical for vascular development and maturation [8], [9]. The hemodynamic forces exerted by blood flow play an essential role in endothelial mechanobiology, as endothelial cells are highly responsive to these forces. Flow influences endothelial cell morphology, gene expression, and survival, with fluid shear stress playing an important role in preventing endothelial apoptosis [6]. Furthermore, disturbed flow patterns are associated with conditions such as vascular inflammation and atherosclerosis [10]. Blood flow is also critical for driving vascular remodeling and ensuring proper vascular development. Studies in chick [11], rat, and mouse embryos [12], [13] demonstrate that the initiation of the first heartbeat and subsequent blood flow are required for vascular structural maturation and, consequently, normal vascular function. However, observational limitations at the time prevented a detailed understanding of how vascular structures remodel under flow. Advances in imaging technology, particularly high-resolution time-lapse imaging, have since revealed that blood flow organizes a hierarchical vascular network by pruning the primitive dense vascular plexus [1], [8]. For instance, these studies demonstrated how blood flow directs the migration of vascular endothelial cells from low-flow regions to areas of higher flow, a process critical for the remodeling and maturation of the vascular network [8], [14]. While these findings underscore the significance of flow-induced remodeling in forming a hierarchical vascular network, to which degree flow-induced remodeling is impacted by VEGF is unclear.

Flow mechanics and VEGF biochemistry are closely linked in vascular remodeling. Flow-induced shear stress activates transcription factors that promote endothelial quiescence and maturation, primarily by downregulating VEGF signaling. This downregulation helps to maintain vessels in a dilated, perfused state, preventing thrombosis and preserving vascular function and stability [6]. In retinal vasculatures, studies have shown that densely branched networks that mimic tumor-like vessels lead to reduced blood flow and tissue hypoxia, as indicated by upregulated VEGF in retinal tissue. In contrast, sparsely branched, normalized vessels provide increased blood supply and oxygenation, resulting in downregulated VEGF levels [15]. More recently, research has shown that blood flow and VEGF signaling together regulate the polarity of vascular cells along the axis of vessel growth in the developing retina, influencing the transition from angiogenic expansion to vascular remodeling [16]. These findings highlight the delicate balance between flow mechanics and growth factor signaling in coordinating vessel maturation. However, the exact balance required for physiological remodeling remains unclear. In addition, animal models have limitations in real-time imaging and flow measurement, making it difficult to accurately study the dynamics of vascular remodeling under varying flow and VEGF conditions. The complexity of whole organisms and the diverse responses to genetic manipulation or surgical intervention make it difficult to isolate the specific effects of altered vascular morphology on blood flow. This highlights the need for in-vitro human vasculature-on-chip platforms to directly assess vascular responses to fluid flow.

Vascularization remains a major bottleneck in the advancing field of human organ-on-chip technology. Without vascular integration and functional perfusion, tissue and organ constructs cannot exceed sub-mm size limits or achieve complex physiological relevance. Vascular network engineering requires a deep understanding of the biophysical relationship between blood flow and network morphology, as well as the dynamic biological processes that drive vessel formation and maturation through remodeling. Despite significant advances over the past decade [17], many human vascular on-chip models remain limited in flow, either being purely static or lacking sustained perfusion. Current models often struggle to maintain long-term perfusion at controlled flow rates and lack well-defined metrics to assess vessel maturation, particularly with respect to flow-induced remodeling. In addition, limitations in the integration of live imaging further hinder the ability to assess the dynamics of vessel maturation and remodeling. While in-vivo studies highlight the need for a balance between flow mechanics and growth factor signaling in vascular remodeling, no in-vitro model has yet successfully replicated this dynamic balance to fully support the maturation of engineered vessels.

Defective vascular remodeling in high-VEGF environments plays a key role in disease progression, with tumor vascularization being a critical example of its pathological impact [1], [2], [18]. Tumor growth beyond a few millimeters and metastasis relies on the development of a vascular network, driven primarily by VEGFs released by tumor cells, which stimulate new vessel formation from the existing host vasculature [19], [20]. Anti-VEGF agents, originally developed to inhibit angiogenesis and restrict tumor growth [19], have shown limited success as monotherapies [21], [22]. However, their efficacy improves when combined with chemotherapy or immunotherapy, likely due to a mechanism known as vascular normalization [21]. Excessive VEGF release by tumor cells leads to an abnormal, tortuous, and dysfunctional tumor vasculature. The concept of vascular normalization suggests that anti-VEGF agents restore the balance between pro- and anti-angiogenic signals within the tumor microenvironment. This balance stabilizes the vasculature and improves its functionality, thereby enhancing drug delivery to the tumor [21]. Nevertheless, combination therapies that aim to exploit vascular normalization to enhance chemotherapy delivery show significant variability in patient outcomes [23], [24]. This variability can be attributed to the critical influence of VEGF levels on tumor behavior, as angiogenesis is highly sensitive to local VEGF fluctuations, especially during vascular maturation [25]. Accurate assessment of tumor vascularity and local VEGF levels during anti-VEGF treatment remains a challenge and may contribute to the inconsistent clinical outcomes observed with these therapies.

Tumor-on-chip models have become valuable tools for studying the tumor microenvironment, largely due to the imaging and real-time analysis limitations of human tumor-bearing animal models. Early efforts to mimic tumor vascularization focused on the formation of non-perfusible, three-dimensional capillary-like structures around tumor spheroids [26]. More recent advances have enabled tumor spheroid invasion by perfusable vascular sprouts, allowing for dynamic interactions within the tumor microenvironment [27]. These perfusable tumor-on-chip models have provided important insights, demonstrating the effects of engineered blood vessels on tumor cell migration [28], immune cell interactions [29], and chemotherapy response [27], primarily under static or limited perfusion conditions. Despite these advances, current models are limited in their ability to provide continuous, controlled perfusion under tumor-level VEGF conditions and to test anti-VEGF drugs or study vascular remodeling under flow. We found that a simpler system containing only blood vessels on the chip - a key component of the tumor microenvironment - provides a powerful approach to overcome these limitations. By focusing solely on the vasculature, this system allows detailed investigation of the effects of flow and VEGF on vascular remodeling, laying the groundwork for a deeper understanding of the tumor microenvironment in future studies.

In this study, we address the limitations of current on-chip vasculature models by investigating how vascular networks respond to long-term flow at both low and high VEGF levels using high-precision, low-pressure syringe pumps for continuous flow control. This approach allowed us to dissect the effect of flow on maintaining the on-chip vascular network phenotype through vascular remodeling and to assess the effect of VEGF levels on this process. By fine-tuning growth factor concentrations, we optimized microfluidic culture conditions that promote the development of perfusable vessels, creating an advanced platform for exploring the intricate relationship between VEGF levels and flow conditions. In high-VEGF environments, we found that anti-VEGF treatment effectively blocks aberrant vascular overgrowth and restores flow-induced remodeling, supporting theories of tumor vessel normalization. In contrast, VEGF neutralization had no effect on remodeling in low-VEGF environments, underscoring the context-dependent efficacy of anti-VEGF therapies and the critical role of the local VEGF environment and tumor vascularity. These findings highlight the need for personalized treatment strategies to optimize therapeutic outcomes. This research lays the groundwork for future real-time analysis of vascular responses to therapies using on-chip models of the human vasculature and opens the door to subsequently explore more complex tumor microenvironments, providing valuable insights to improve clinical outcomes of anti-angiogenic treatments.

## Results

### Enhancing vascular connectivity and perfusability through optimized culture conditions and chip design

To study the effects of flow and growth factors on vascular remodeling, it is essential to recapitulate the process of vessel development in a durable microfluidic chip environment that allows for the perfusion of engineered vascular networks. Since vessel formation and maturation occur over days to weeks, we adapted a polydimethylsiloxane (PDMS)-based microfluidic culture system to be compatible with precision syringe pumps, live imaging, and maintainability for up to 12 days. We used a 5-channel PDMS chip configuration, with the vascular compartment centrally located and separated from the two media channels on either side by micro-posts. To ensure uniform expansion of the vascular network, the vascular compartment is flanked by two additional stromal cell chambers (Fig. 1A). Stromal cells, such as fibroblasts, are known to secrete factors that promote vessel morphogenesis, enhance stability [30], [31], and facilitate the formation of open, perfusable lumens on-chip [32]. The separation of vascular and stromal cells serves two main purposes. First, it allows quantitative analysis of vessel formation and remodeling using vascular tubes composed only of endothelial cells, mimicking the conditions observed in many aggressive tumors. Second, it streamlines the extraction of vascular network structures directly from brightfield image segmentation of vessel walls, providing an efficient approach for real-time network analysis. We used a well-established technique [32] in which micro-posts laterally separate the vascular chamber from the adjacent media channels. These posts not only contain the gel during injection but also support the formation of perfusable luminal openings, as the vascular tubes tend to remain open at the gel-liquid medium interface toward the stromal cell chambers at the inter-post openings (Fig. 1A). When endothelial cells suspended in fibrin hydrogel are injected into the central vascular chamber, the cells undergo vasculogenesis within approximately six days (Figs. 1A, F).

**Figure 1.**
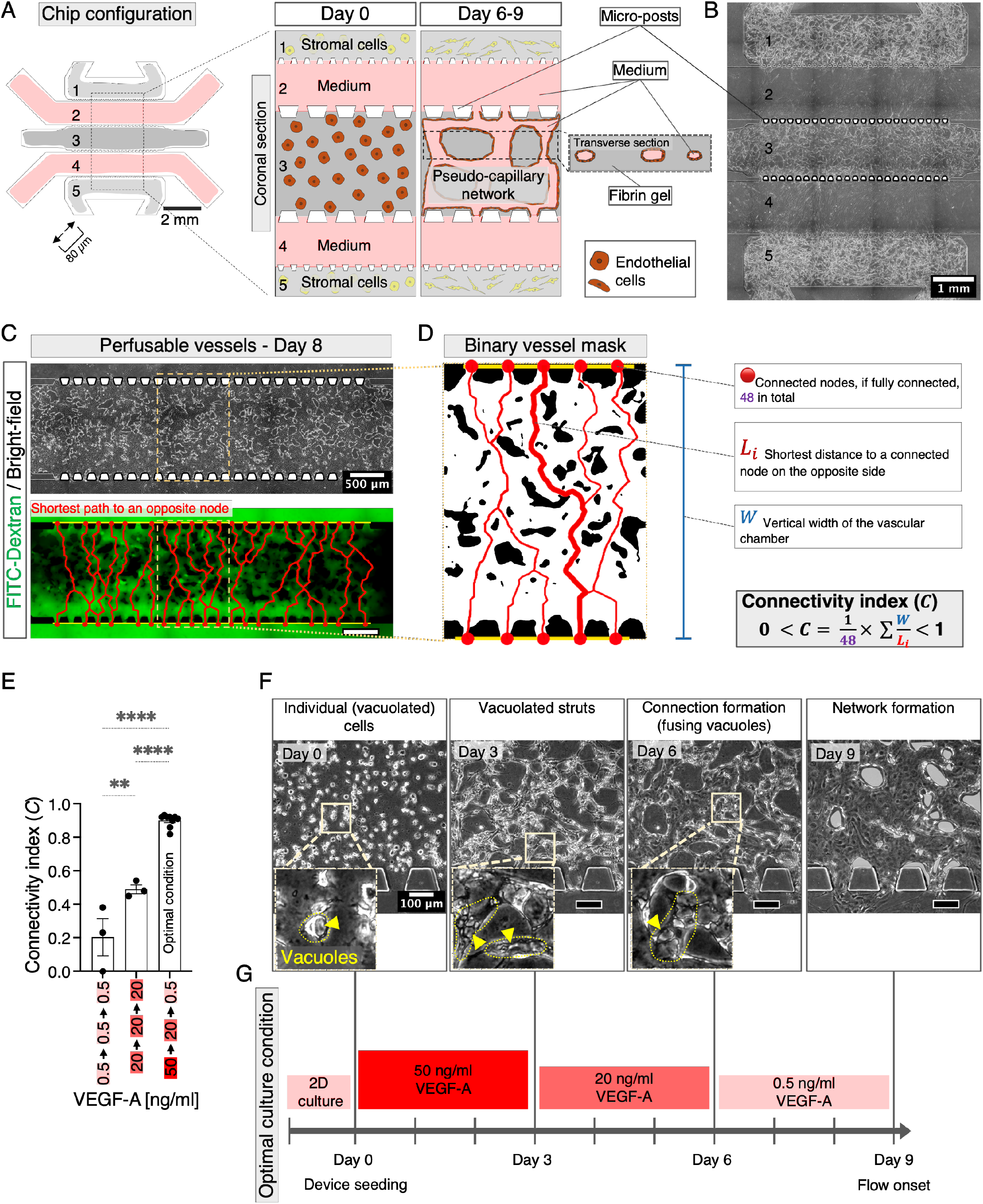
Microfluidic chip design and optimization of culture conditions for perfusable and connected vessels. **(A)** The microfluidic chip design features five channels, with the central vascular chamber flanked by two stromal cell chambers. The vascular chamber is separated from the media channels by micro-posts, with a chip height of 80 µm. **(B)** Brightfield image of the chip showing an interconnected vascular network in chamber 3, separated from the media channels by micro-posts, indicated by white trapezoids. **(C)** Brightfield image of the on-chip vascular network extending over an area of 5.3 x 1.3 mm between micro-posts, perfused with FITC-Dextran. Red lines indicate the shortest vascular connections between nodes. **(D)** Schematic of how the connectivity index *C* is determined using a black and white vessel mask segmented from the region shown in (C). Red circles indicate connected vessel openings along the upper and lower yellow lines marking the gel-media channel interface. Red paths (''*Li*'') represent the shortest path between oppositely connected nodes within the network skeleton. **(E)** Quantification of the connectivity index *C* across three culture conditions on day 9, with VEGF-A concentrations (ng/ml) on the horizontal axis and media changes occurring every three days from day 0 to day 9. Data represent mean ± S.E.M. One-way ANOVA with Tukey's multiple comparison test was performed; **p < 0.01; ****p < 0.0001 (n ≥ 3). **(F)** Brightfield images of endothelial cells undergoing tubulogenesis on-chip. Arrowheads point to vacuoles within cells, within elongated struts, and within fusing tubes (outlined with yellow dotted lines). White-shaded areas indicate avascular portions of the gel (''gaps within the vascular network'') on day 9. **(G)** Timeline of sequential growth factor concentration adjustments from day 0 to day 9, with the maximum connectivity index *C* indicating the optimal culture condition. Images are representative of at least three independent samples.

To enable precise monitoring and assessment of vascular tube formation and remodeling dynamics, we designed PDMS chips with a height of 80 µm. This reduced height minimizes three-dimensional structural complexity and promotes planar vessel growth predominantly in the x-y plane within chamber dimensions of about 5 × 1.3 mm. By aligning the centerlines of the endothelial tubes near the mid-height of the chamber (Figs. 1A, B), this design enhances network visibility within a single focal plane, facilitating real-time brightfield imaging and two-dimensional network quantification. The planar morphology not only ensures accurate and reliable data acquisition but also preserves the hierarchy of vessel diameters within the network, making it ideal for studying the dynamics of flow-induced remodeling. To improve the durability of our chips, we added aprotinin to the culture medium. As fibrin is prone to proteolytic degradation, aprotinin, as a potent protease inhibitor, is commonly used to prevent fibrin degradation and maintain gel stability. Thus, aprotinin helps to create a durable scaffold for cell adhesion and growth while allowing cells to remodel and migrate by secreting matrix metalloproteinases [33]. Monitoring of the engineered vessels from day 0 to day 12 showed that the conventional protocol of adding aprotinin only to the gel mixture does not ensure long-term durability. Real-time observations showed gel degradation in approximately 40% of the chips, likely due to diffusion of aprotinin (6.5 kDa) out of the fibrin gel and into the culture medium [34]. We found that the addition of aprotinin to the culture medium was necessary to stabilize the gel integrity and potentially better preserve its mechanical properties.

To establish a culture protocol that supports the recapitulation of physiological vascular development through the two main phases of vasculogenic network formation and subsequent angio-adaptive network maturation, we used real-time observation of the developing networks. This was facilitated by an incubator microscope and the use of chambered microscope slides featuring walls and a lid to house the PDMS chips. This setup ensured optimal control of the culture environment, particularly with respect to medium evaporation, which is critical for the stability of the forming vascular structures within the chips. Our live observations revealed a characteristic three-stage sequence of events leading to the formation of perfusable and interconnected vascular networks within our chips. These stages included: 1) An initial vasculogenic step where individual cells vacuolate to form lumens and rearrange into a network of elongated struts/rods, giving rise to the primitive network structure (Fig. 1F-Day 0, Day 3). 2) A phase of connection formation during which vacuolated struts fuse end-to-end and begin to form interconnected networks. By the end of this stage, most lumenized structures are interconnected, although some still contain separate vacuoles without continuous lumens (Fig. 1F-Day 6). 3) An early stage of network maturation when all lumens are fully interconnected, and more than 70% of the network can be infused with a fluorescent tracer (Figs. 1C, F-Day 9).

Inspired by the critical role of VEGF signaling in vascular development and its regulation as a key step in stabilizing and maturing vessels [6], [16], we used a stepwise approach to adjust VEGF concentration. From day 3 to day 6, we gradually reduced VEGF levels in the endothelial growth medium. This reduction was further extended from day 6 to day 9 to induce vessel maturation. This sequential reduction facilitated the transition from active sprouting and growth to stabilization and maintenance of the vascular network. We tested four different protocols with different VEGF levels in the growth medium, with and without sequestering of VEGF, to find an optimal culture condition for prolonged perfusion (Fig. 1E). We scored culture condition with a connectivity index *C*, which measures both the interconnectivity of the vessels and their connection to the medium channel through lumen openings at the inter-post openings (Fig. 1D). Chips with vascular networks of connectivity index *C* of 0.7 or larger were experimentally determined to be optimal since networks of lower connectivity were prone to rupture under perfusion. The optimal VEGF condition started with an “ultra-high” VEGF concentration of 50 ng/ml, recognized in the literature as an “pro-angiogenic medium” [35], and was gradually reduced to a “high” level of 20 ng/ml on day 3, followed by a further reduction to a “baseline” level of 0.5 ng/ml (low-VEGF) on day 6. The baseline level was then maintained until day 12. The baseline VEGF level of 0.5 ng/ml corresponds to the typical range found in the blood and muscle tissues of healthy individuals (reported as 0.3 ng/mL in whole blood and 0.16-0.48 ng/mL in the extracellular space of muscle tissue [36]). This sequential reduction (50-20-0.5 ng/ml, Fig. 1G) maximized the connectivity index *C*, with an average connectivity of 0.86 (Fig. 1E).

### Optimizing chip geometry and inflow rate is essential for long-term perfusion

In our experiments, poorly connected vessels often ruptured when subjected to continuous flow conditioning using fixed flow rate syringe pumps. Our real-time imaging showed that vessel rupture was characterized by detachment of endothelial cells from the vessel walls, followed by their migration away from the rupture site and adoption of a two-dimensional phenotype (Fig. 2B). In contrast, brightfield images of intact, unruptured flow-conditioned networks showed vessel walls that responded dynamically to flow without disassembling (Fig. 2A). Post-perfusion immunostaining further confirmed the structural differences between ruptured and unruptured vessels. In ruptured vessels, the endothelial cells adopted a two-dimensional phenotype, as shown by cross-sectional images marked by the luminal marker ICAM-2 and the basement membrane marker collagen IV (Fig. 2D). Conversely, unruptured vessels retained a three-dimensional hollow lumen, indicating tubular vascular architecture (Fig. 2C). To better understand the relationship between shear stress and vessel rupture, we performed computational fluid dynamics simulations using brightfield images of the vascular networks on day 9, when perfusion began. By comparing ruptured networks with intact ones, we found that endothelial tubes were prone to rupture when exposed to maximum shear stress above 0.05 Pa (0.5 dyn/cm2, Fig. 2F). In contrast, when the maximum shear stress remained between 0.01 and 0.05 Pa (Fig. 2E), the vessels underwent dynamic remodeling under flow, characterized by the repositioning of the vessel walls while remaining structurally intact (Fig. 2C). These observations highlight the critical role of shear stress for vessel stability during perfusion.

**Figure 2.**
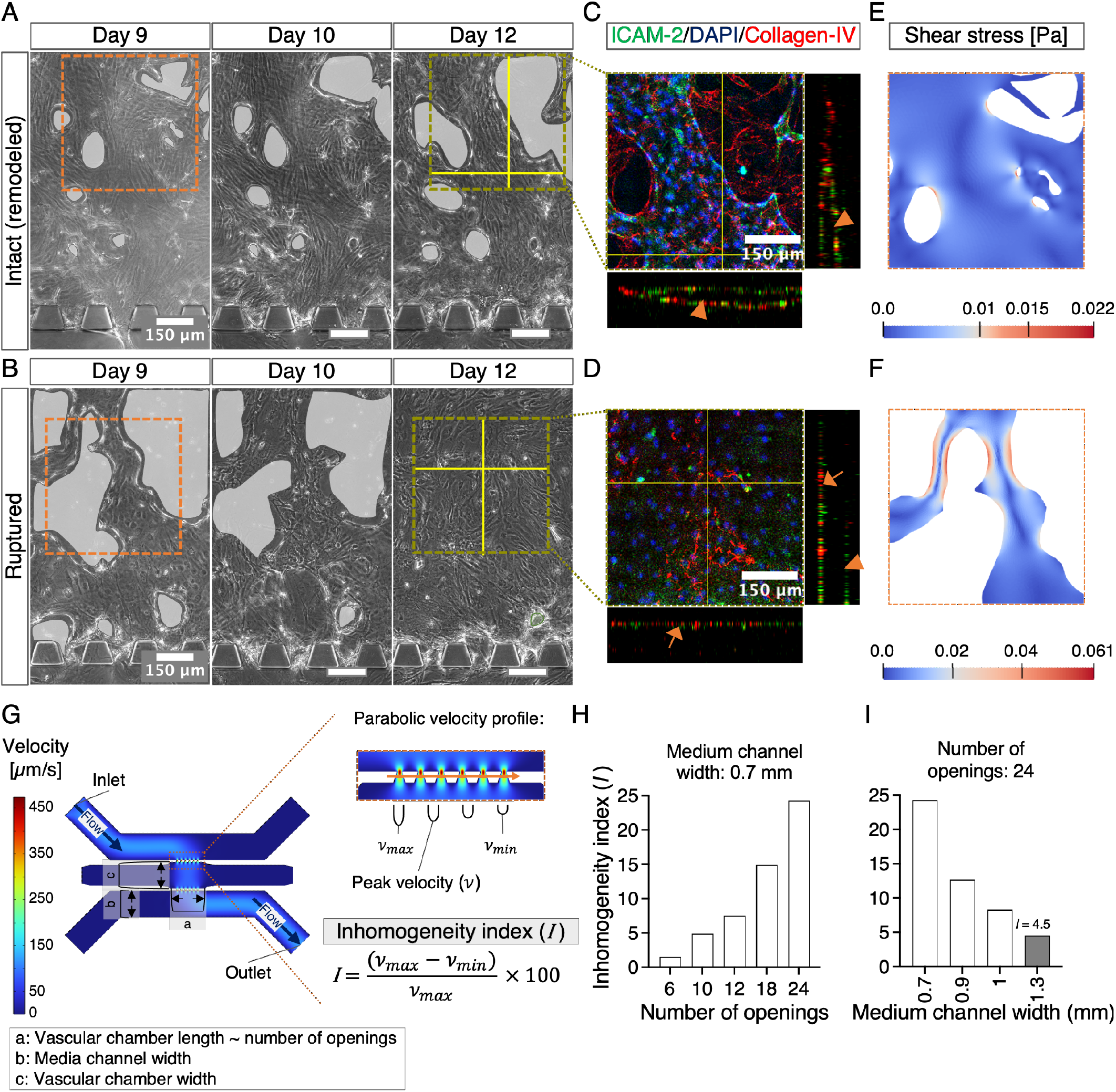
Optimization of chip geometry and flow conditions for long-term perfusion. **(A)** Brightfield images show a selected region of the vessel network at the start of flow conditioning on day 9 and on days 10 and 12. White-shaded areas highlight gaps in the network, indicating the position of the vessel walls. The images show that the vessel walls rearrange under flow but remain intact throughout this period. **(B)** Brightfield images of a selected region of another flow-conditioned network from day 9 to day 12 show white-shaded gaps that mark the position of the vessel walls on day 9 and day 10. The vessel walls show signs of collapse, and the gaps are later filled by two-dimensional cell layers, indicating vessel rupture. **(C)** Immunostaining on day 12 for ICAM-2 (green), Collagen-IV (red), and DAPI (blue) shows hollow, three-dimensional vessels (arrowheads) in unruptured samples. **(D)** In contrast, ruptured vessels exhibit a monolayer endothelial morphology (arrows) with remnants of three-dimensional structures (arrowhead). **(E)** Simulated shear stress visualized in the indicated region of the unruptured vessels in (A)-day 9. **(F)** Simulated shear stress visualized in the indicated region of the ruptured vessels in (B)-day 9. **(G)** Simulated flow velocity in an empty chip geometry showing spatially varying velocity profiles across the openings between the media channel and the vascular chamber. The difference between the peak velocities is quantified as the inhomogeneity index *I*. **(H)** Inhomogeneity index *I* calculated from simulated velocities in chips with different numbers of openings, where the media channel width is fixed at 0.7 mm, and the vascular chamber width is fixed at 1.3 mm. **(I)** Inhomogeneity index *I* calculated from simulated velocities in chips with different media channel widths, with the number of openings fixed at 24 and the vascular chamber width fixed at 1.3 mm.

Next, we investigated the effect of chip geometry and inflow rate on vessel shear stress. Since vessel shear stress is directly proportional to the flow rate of the individual vessel, we investigated how chip geometry and inflow rate affect this parameter. Averaging over an entire microfluidic chip, we estimated the typical vessel flow rate as the chip inflow rate divided by the number of connections between the top and bottom openings. To quantify these connections, we used the connectivity index *C*, which measures the fraction of functional connections between openings. The number of connections was estimated as the product of the connectivity index *C* and the total number of openings. With our optimized culture protocols, connectivity indices averaged around 0.9 and consistently exceeded 0.7 across samples (Fig. 1E). Once the culture conditions were optimized to maximize the connectivity index *C*, the remaining variable under our control was chip inflow rate. Based on reported blood velocity values in the human microvasculature [37], we approximated flow rates of 0.01 to 0.1 µl/min for 50 µm vessels. Scaling these values to our chip configuration of 24 fully connected apertures, we calculated a chip flow rate range of approximately 0.24 to 2.4 µl/min. From this range, we selected 1 µl/min as our initial test flow rate. However, at µl/min, poorly connected networks exhibited vessel rupture (Fig. 2B). To mitigate this, we reduced the flow rate to 0.5 µl/min, ensuring that the shear stress remained within a tolerable range, particularly for networks with a minimum connectivity index *C* of 0.7.

These observations highlighted the importance of not only controlling the typical vessel shear stress but also minimizing shear stress variations across the network. Increasing the number of openings from the typical range of 3-10 used in previous studies [32], [38] to 24 openings significantly reduced the statistical risk of poor connectivity. This adjustment reduced the impact of a single missing lumen connection at an opening on overall network connectivity and reduced the likelihood of localized areas of high shear stress that could lead to vessel rupture. Thus, the balance between chip inflow rate, connectivity, and number of openings is critical to support both functional maturation and mechanical stability of vascular networks during long-term perfusion.

Recognizing the importance of variations in flow shear stress, we investigated how specific aspects of chip design contribute to achieving homogeneous flow. The chip design includes three main elements: the number of openings (which also determines the length of the media channels and the vessel chamber), the width of the media channels, and the width of the vessel chamber (Fig. 2G). To evaluate their effect on flow homogeneity, we performed COMSOL simulations using a two-dimensional model of our chip geometry (Fig. 2G). The simulations were performed with an inflow velocity of 80.13 µm/s, corresponding to the experimental volumetric flow rate of 0.5 µl/min, an inlet width of 1.3 mm, and a chip height of 80 µm. The inlet was positioned at the top left corner of the chip, with a symmetrical outlet at the bottom right corner. The fluid flow enters the media channel at the inlet and moves from left to right, gradually flowing into the vessel chamber through the openings between the upper media channel and the vessel chamber. As the fluid moves further along the media channel, it experiences increasing friction along the channel walls, resulting in a gradual decrease in flow velocity at the openings from left to right. To quantify this effect, we introduced an inhomogeneity index *I*, which measures the percentage difference between the highest and lowest local maxima in the velocity profile (peak velocity) along a line passing through the center of the micro-post row (Fig. 2G). A higher inhomogeneity index *I* indicates a more uneven flow distribution, while an index of zero reflects a completely uniform flow. Using the inhomogeneity index *I* as a benchmark, we next evaluated how variations in chip geometry influence flow distribution and homogeneity.

We first examined how increasing the number of openings between the media channels and the vascular chamber affected flow homogeneity. Increasing the number of openings offered two key advantages: (a) it statistically reduced the risk of poor connectivity by providing more opportunities to form connected, open vessel lumens, as discussed previously, and (b) it increased the interface between the media channels and the vascular chamber, thereby improving nutrient diffusion into the gel. To take advantage of these benefits, we increased the number of openings from the typical range of 3-10 used in previous studies [32], [38] to 24 openings. However, this increase in openings also increased the length of the media channels, which increased the friction for the fluid moving from the inlet to the rightmost opening. Consequently, increasing the number of openings from 6 to 24 resulted in a 16-fold increase in the inhomogeneity index *I*, from 1.5% to 24.3% (Fig. 2H). While this increase in flow inhomogeneity was suboptimal, it was a necessary trade-off to improve the perfusability of our chips and could be mitigated by optimizing other design factors, as discussed below.

To address the increased flow inhomogeneity caused by the increased number of openings, we identified media channel width as a key design parameter. While increasing the length of the media channel increases the overall flow friction, widening the media channel can counteract this effect by reducing the flow resistance at a comparable rate. Our simulations showed that increasing the media channel width from 0.7 mm [32] to 1.3 mm (the maximum allowed by the PDMS housing slides) significantly reduced the inhomogeneity index *I* by approximately 5.4 times, from *I* = 24.3% to *I* = 4.5% (Fig. 2I). This adjustment effectively mitigated the adverse effects of a greater number of orifices on flow inhomogeneity, resulting in a more uniform flow distribution throughout the vascular network. Although some degree of inhomogeneity is inevitable in our parallel channel configuration, we found that an inhomogeneity index *I* of 4.5% was acceptable for our system.

Next, we investigated the effect of chamber width on flow homogeneity. This width is critical for nutrient delivery to the cells, which relies on the diffusion of molecules from the media channels into the gel within the vascular chamber during the early vasculogenic phase (day 0 to day 3). It particularly affects the vascular connectivity in the center of the chamber. Based on widths reported in the literature (1 to 4 mm) [32], [38], [39] and the constraints of our PDMS housing slides (1 to 2.5 mm), we chose a width of 1.3 mm. This choice balanced improved vessel connectivity with a chamber size large enough to support vascular network expansion for flow-induced remodeling studies. Simulations showed that increasing the width from 1 mm to 2.5 mm slightly increased the inhomogeneity index *I* by 1.16-fold (from *I* = 4.3% to *I* = 5%, Supplementary Fig. S1). Notably, widening from 1 mm to 1.3 mm resulted in only a 1.04-fold increase (to *I* = 4.5%), after which the index plateaued at 2 mm. These results demonstrate that variations in chamber width within our range had negligible effects on flow inhomogeneity, making 1.3 mm an optimal balance for connectivity and network development.

Balancing flow homogeneity, nutrient diffusion, and vessel connectivity was critical to our chip design. While optimizing flow homogeneity was essential, we also aimed to leverage the positive effect of increasing the number of openings to increase the connectivity index *C*, while at the same time addressing the limitations of nutrient diffusion. By combining a narrower vascular chamber with an expanded media-vessel interface, i.e., a greater number of openings, we improved nutrient delivery and maximized the connectivity index *C*. Taking all these factors into account, our optimized chip design includes 24 openings, a media channel width of 1.3 mm, and a vascular chamber width of mm. This design achieved a minimum connectivity index *C* of 0.7 and ensured that the shear stress across the flow-conditioned samples remained between 0 and a maximum of 0.05 Pa. This balance of factors successfully prevented vessel rupture during subsequent perfusion experiments and provided a stable platform for long-term perfusion of the vascular network.

### Flow-driven vascular remodeling by pruning preserves on-chip network phenotype and increases flow velocities

After optimizing culture conditions and chip geometry, we first examined network maturation under static conditions, with VEGF reduced to a baseline level of 0.5 ng/ml from day 6 onward (Fig. 3A). From day 9 to day 12, the static vessels exhibited gradual continuous growth, leading to the lateral fusion of branches (Fig. 3B and Supplementary Movie S1). This persistent growth, even after the reduction of VEGF, suggested that mechanical cues from flow might be essential for maintaining the network-like structure of the vessels and preventing a loss of hierarchy due to persistent growth. Based on these findings, we proceeded to perfuse the chips in the next experiment phase.

**Figure 3.**
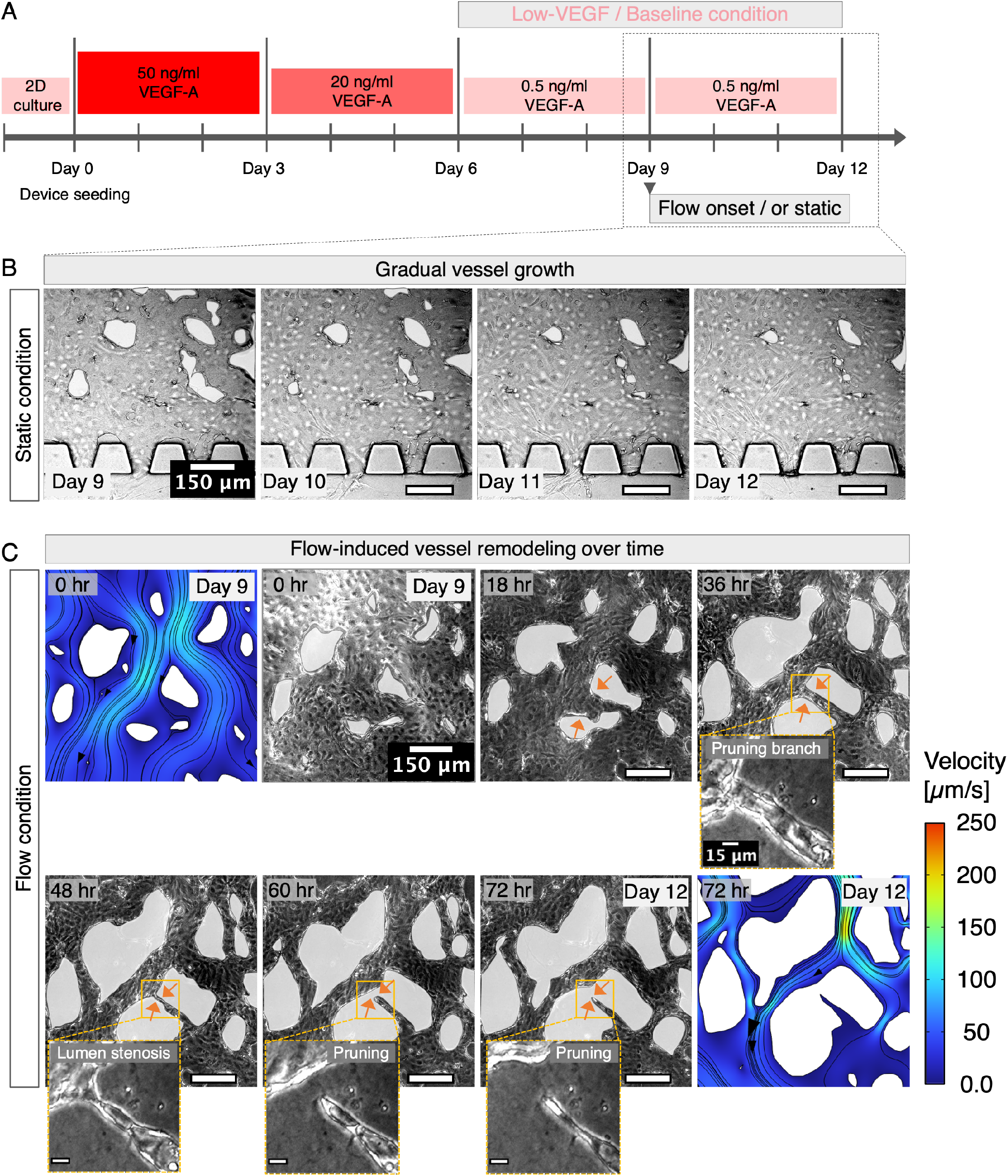
Flow conditioning induces network remodeling via vessel pruning. **(A)** Timeline of culture conditions from day 0 to day 12, with flow conditioning under low/baseline VEGF levels from day 9 to day 12. **(B)** Brightfield images of a selected region in a static network from day 9 to day 12 show continuous, gradual growth, as indicated by the decreasing size of the white-shaded avascular gaps over time. **(C)** Brightfield images of a selected region in a flow-conditioned network, with avascular gaps white-shaded, the borders of which indicate vessel walls. Arrows in the 18-hour image indicate wall rearrangement leading to lumen narrowing. Magnified views of the 36- and 48-hour images highlight lumen stenosis and breakage at a pruning branch (arrows), while arrows in the 60- and 72-hour images show pruning branch retraction. Simulated velocity fields from the 0- and 72-hour (day 9 and day 12) images show velocity magnitude, flow streamlines, and flow direction before and after remodeling. Brightfield images and simulated fields represent at least three independent samples.

To understand the effect of flow on vascular remodeling, microfluidic chips were perfused with a low-pressure syringe pump system allowing precise sub-nanoliter fluid dosing. Each microfluidic chip was paired with a pump module equipped with a glass syringe and a valve with a pressure sensor. The flow rate for each pump module was consistently maintained at 0.5 µl/min using the manufacturer's software. Live imaging was performed at 2-6 hours intervals to monitor the networks and to ensure that no rupture occurred throughout the three-day perfusion experiment. We employed images to analyze the structural remodeling of the vascular networks that occurs through the rearrangement of endothelial cells upon perfusion. Our engineered vasculature showed stepwise regressive vascular remodeling (“pruning”) described as vascular anastomosis in reverse in animal models [40] involving lumen stenosis and breakage followed by endothelial cell retraction (Fig. 3C and Supplementary Movie S1).

To assess flow-modulated structural changes in the flow-conditioned vessels, we compared the vascular networks before and after perfusion (“pre-” and “post-mask”). Vessel masks were extracted from brightfield images of the entire vascular chamber using a machine-learning algorithm to segment vessel walls. By superimposing the pre- and post-masks, we identified regressive pruning sites by network branches present in the pre-mask but absent in the post-mask (Fig. 4A and Supplementary Movie S1). Regression under perfusion contrasts with the observation of continuous growth in static samples (Fig. 4A and Supplementary Movie S1). We further performed co-immunostaining for ICAM-2 and Collagen-IV on day 12. This allowed us to identify sites of regressive pruning, which were characterized by empty Collagen-IV sheaths, representing the remaining basement membrane, and the absence of endothelial cells (ICAM-2) [40]. Confocal z-stack images of flow-conditioned samples at sites of regressive remodeling indeed showed empty Collagen-IV remnants near retracting luminal structures surrounded by viable hollow vessels, confirming vessel pruning as the mechanism driving network remodeling under flow conditions (Fig. 4B). In contrast, z-stack images of the co-stained static samples revealed merging tubular structures at sites of lateral vessel fusion (Fig. 4B).

**Figure 4.**
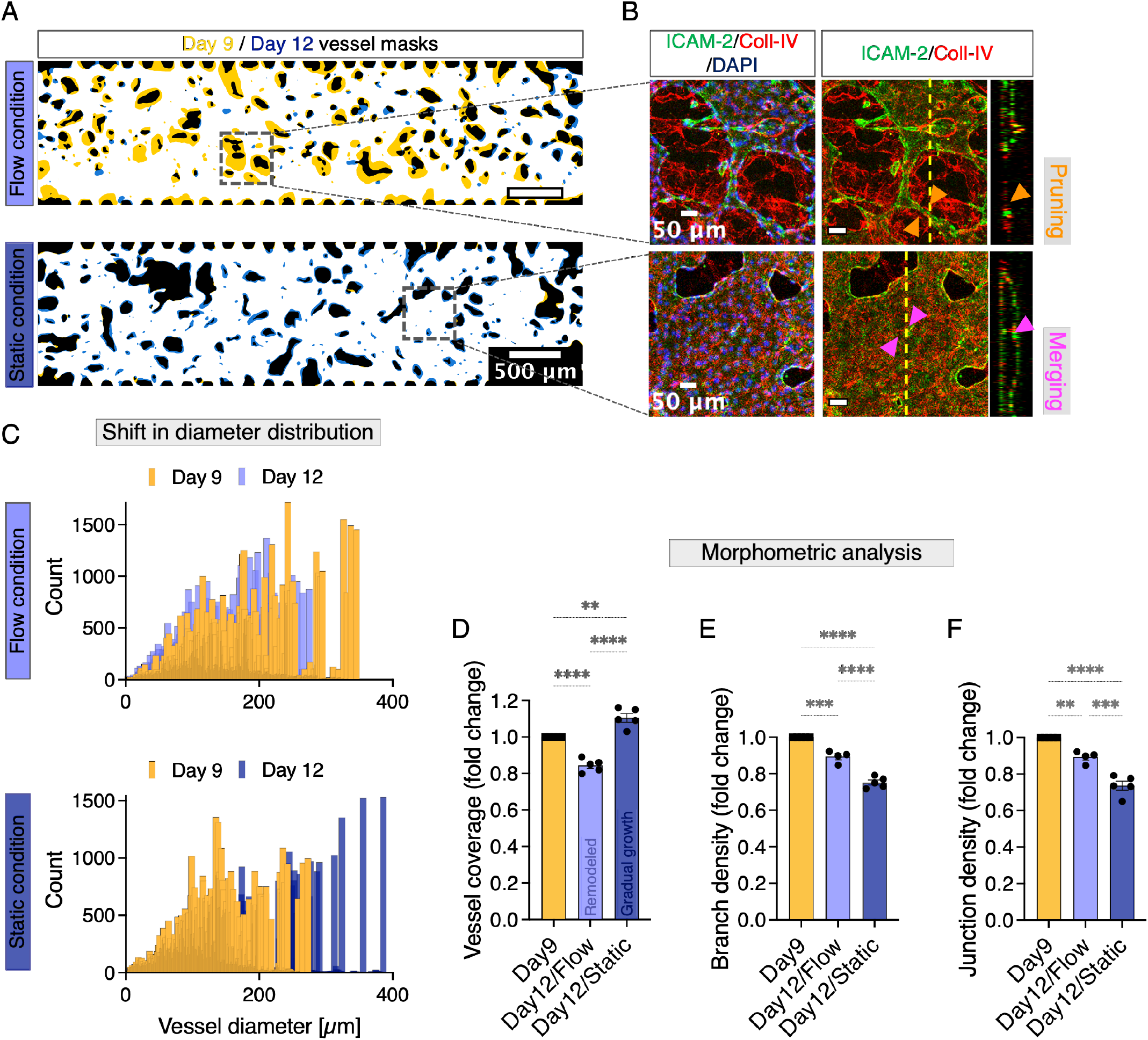
Flow-induced vascular remodeling preserves on-chip network phenotype. **(A)** Superimposed day 9 and day 12 masks showing pruning sites (yellow segments) in the flow-conditioned sample and enlarged vessels (blue segments) in the static sample. **(B)** Immunostaining images of the endothelial lumen labeled with ICAM-2 (green), the basement membrane labeled with Collagen-IV (red), and the nuclei labeled with DAPI (blue) in the indicated regions of the flow conditioned and static samples in (A). The arrowheads in the flow-conditioned sample indicate pruning sites in the two-dimensional and corresponding cross-section images. In the static sample, arrowheads indicate merging vessel walls in the two-dimensional and corresponding cross-section images. **(C)** Vessel diameter distribution on day 9 and day 12 for the same flow-conditioned and static samples as in (A) show broadening in static conditions and narrowing under flow. **(D - F)** Morphometric analysis shows network remodeling under flow compared to loss in network architecture under static conditions: **(D)** Quantification of vessel coverage by area density, **(E)** Vessel branching density as the number of branches per mm of vessel length, and **(F)** Junction density as the number of junctions per mm of vessel length in flow conditioned and static samples on day 12, shown as fold change compared to the respective day 9 values. Data represent mean ± S.E.M. Ordinary One-way ANOVA with Tukey's multiple comparisons test was performed; **p < 0.01; ***p < 0.001; ****p < 0.0001 (n ≥ 4). The vessel masks and fluo-rescent images represent at least three independent samples.

Morphometric analysis using the pre- and post-masks showed that remodeling of the vascular network in flow-conditioned samples led to a shift in vessel diameter distribution toward thinner tubes, while static controls exhibited a thickening of tubes from day 9 to day 12 in line with continuous overgrowth (Fig. 4C). Along with regressive remodeling in the flow-conditioned samples, we observed a decrease in vessel coverage (Fig. 4D), measured by vessel mask area density, and thus network refinement, characterized by a slight reduction in vessel junction and vessel branch density (Figs. 4E, F). The continuous overgrowth of static samples, on the contrary, resulted in a gradual increase in vessel coverage (Fig. 4D) and a large reduction in branching and junction density, driven by lateral fusion of vessel branches leading to network diminishment (Figs. 4E, F).

To evaluate the effect of regressive vascular remodeling on flow distribution in vessels, we simulated flow in two-dimensional pre- and post-masks (day 9 and day 12) of the flow-conditioned samples. As a control, hypothetical flows were also simulated in static samples for the same time points. Visualizations of flow velocity across the networks revealed that continuous outgrowth in static samples from day 9 to day 12 resulted in regions of decreased flow velocity (Fig. 5A), likely due to vessel dilation. In contrast, flow-conditioned samples exhibited regions of increased velocity at day 12 compared to day 9 (Fig. 5A) due to vessel constriction and network reorganization. To enable a meaningful comparison of velocity distributions across networks with different vessel coverage, we normalized the velocities within each network using the Darcy velocity. The Darcy velocity, a standard metric for flow in porous media, represents a reference velocity defined by the inflow rate and the cross-sectional area of the network available for perfusion. Superimposing the normalized velocity distributions from days 9 and 12 revealed a consistent trend: flow-conditioned samples shifted toward higher velocities, whereas static samples shifted toward lower velocities (Figs. 5B, C). This pattern was observed for all three flow-conditioned samples and the three static controls (Supplementary Fig. S2). The same trend was evident in the raw velocity data before normalization, with static samples showing a shift toward lower velocities and flow-conditioned samples showing a shift toward higher velocities (Fig. 5D).

**Figure 5.**
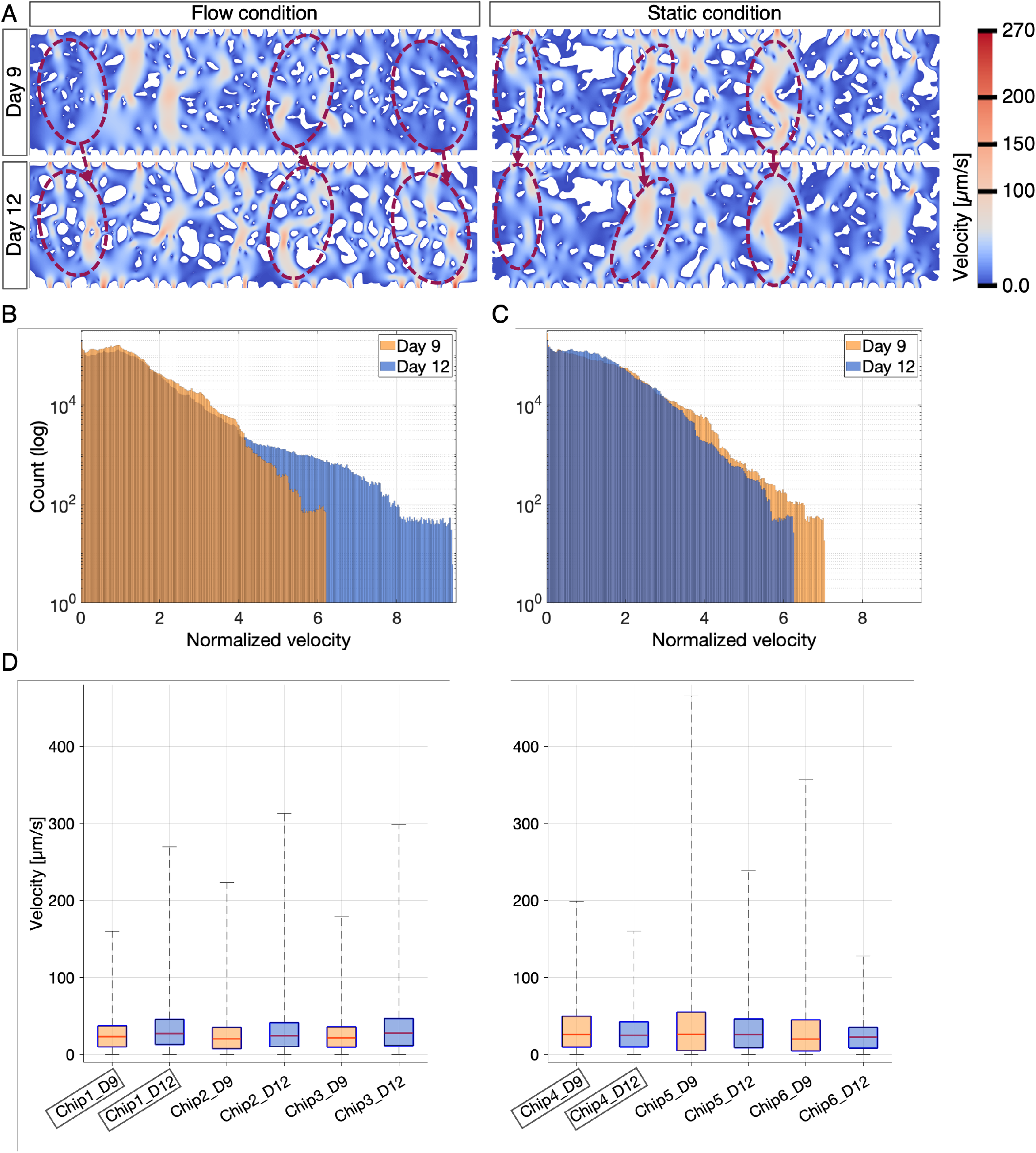
Flow-induced remodeling of vascular networks shifts the velocity profile toward higher flow velocities. **(A)** Simulated flow velocities within the two-dimensional vessel masks for the flow-conditioned and static samples shown in Figure 3A at the onset of flow on day 9 and again on day 12. Under flow conditioning, regions with increased local velocities on day 12 after perfusion compared to before perfusion on day 9 are high-lighted. In the static sample, regions with decreased local velocities on day 12 compared to day 9 are highlighted. **(B)** Velocity distribution for the flow-conditioned and **(C)** static samples from (A), normalized by the Darcy velocity calculated for each vascular network, show a shift to higher flow velocities under flow and, contrary, to lower flow velocities under static conditions. Simulated velocity fields and the plotted velocity distributions represent at least three samples per group. **(D)** The distribution of velocity values before normalization is shown for three independent flow-conditioned and static samples on day 9 (D9) and day 12 (D12). The chips highlighted with boxes are the same flow-conditioned and static samples as those in (A), (B), and (C). Note that for the static samples, the visualized velocity fields of day 9 and day 12 are hypothetically modeled since these samples were not exposed to actual flow conditions.

Our results demonstrate that under reduced VEGF following the initial vasculogenic phase, flow is critical for maintaining the network phenotype. The onset of flow prevents continuous, uncontrolled vessel growth, preserves network hierarchy, and induces remodeling, consistent with observations in animal models. This remodeling results in an increase of high-flow regions in the remodeled vascular networks, whereas flow velocities would be reduced under uncontrolled growth of static samples. Enhanced flow velocities suggest improved perfusion, facilitating more efficient delivery of nutrients and removal of waste products. This further highlights the essential role of flow in maintaining the structural and flow properties in our vasculature on a chip.

### VEGF neutralization is key to restoring flow-induced vascular remodeling under tumor-mimicking high-VEGF conditions

Vascular abnormalities in conditions such as tumor growth and metastasis are closely associated with excessive release of VEGF [19], [21]. Since defective vascular remodeling is a hallmark of tumor aggressiveness [2], [24], we aimed to evaluate flow-induced remodeling under high VEGF conditions using our vascular chips. To mimic the tumor microenvironment, we maintained a high VEGF concentration of 20 ng/ml in the culture medium after the initial vasculogenic phase (day 0 to day 3, 50 ng/ml VEGF, Fig. 6B) throughout the subsequent connection formation and early maturation phases (day 3 to day 6 & day 6 to day 9, Fig. 6B). This contrasts with our standard protocol, where VEGF levels are reduced to a baseline of 0.5 ng/ml from day 6 to day 9. The selected VEGF concentration reflects a 40-fold increase in VEGF observed in lung tumor effusions [41] roughly aligning with levels reported in the blood [42] and tumor cytosol [36] of cancer patients.

**Figure 6.**
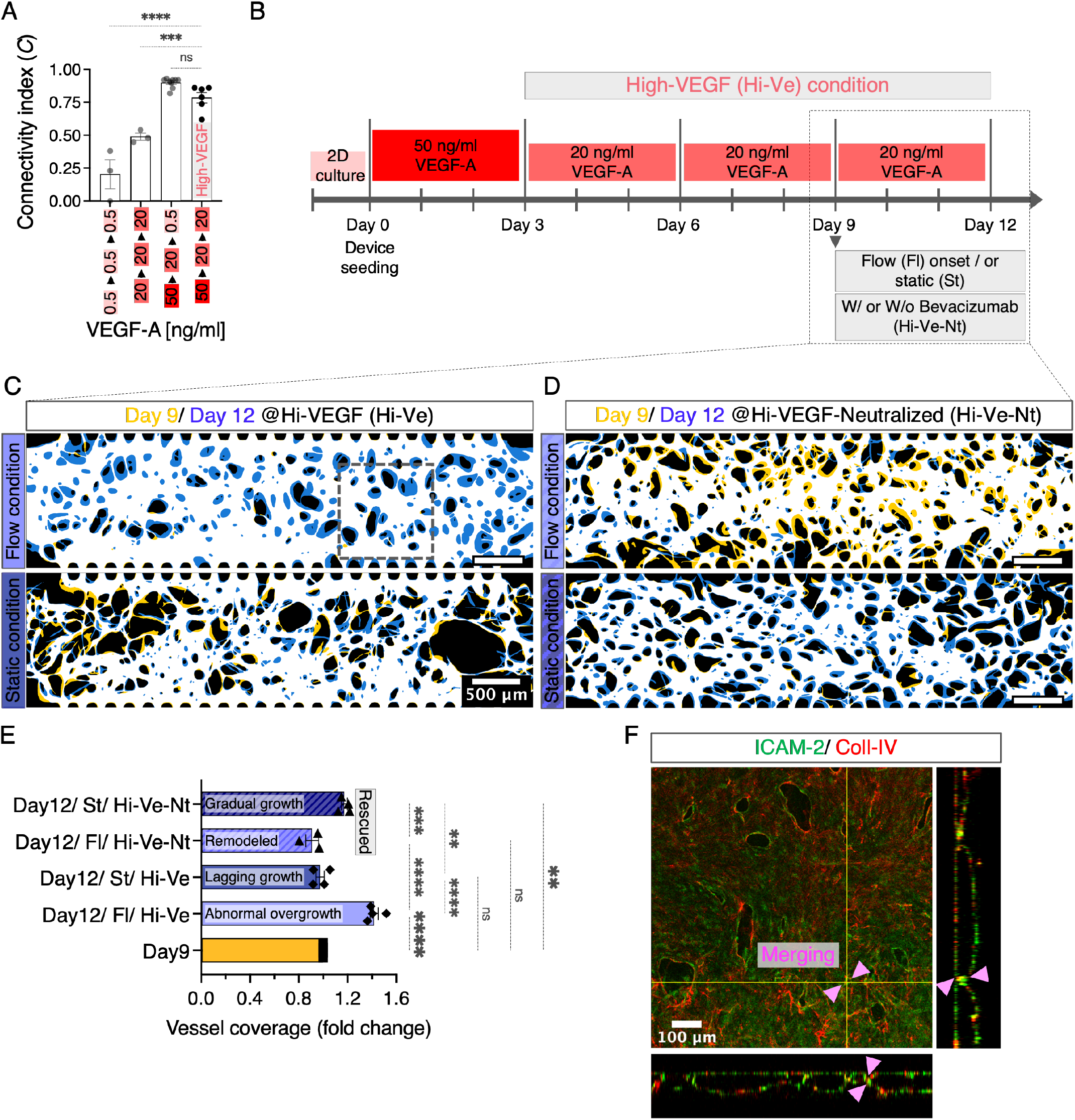
Cancer drug-induced VEGF neutralization prevents abnormal flow-driven vessel growth under high-VEGF conditions and restores vascular remodeling. **(A)** Quantification of the connectivity index *C*, on day 9, under high VEGF (Hi-VEGF) condition in comparison with the previously presented culture conditions (faded, same data points as in Fig. 1E) with VEGF-A concentrations (ng/ml) on the horizontal axis and media changes occurring every three days from day 0 to day 9. **(B)** Timeline of growth factor concentrations from day 0 to day 12, showing the sequential adjustment of VEGF-A levels. **(C)** Superimposed day 9 and day 12 masks of flow-conditioned and static samples under high-VEGF. Flow-conditioned sample shows significant overgrowth with enlarged vessels (blue segments), while the static sample shows delayed vessel growth, with fewer blue segments and some regressed branches (yellow segments). **(D)** Superimposed day 9 and day 12 masks of flow-conditioned and static samples under high-VEGF plus bevacizumab. VEGF neutralization restores the flow-induced remodeling in the flow-conditioned sample indicated by pruning of branches (yellow segments). It also restores the gradual vessel growth in the static sample (blue segments). **(E)** Quantification of vessel coverage by area density in day 12 vessels under high-VEGF (Hi-Ve) and high-VEGF plus bevacizumab (Hi-Ve-Nt) conditions for both flow conditioned (Fl) and static (St) samples, shown as fold change compared to day 9 values. Vascular remodeling is rescued under VEGF neutralization with bevacizumab. **(F)** Immunostaining images of the endothelial lumen labeled with ICAM-2 (green) and the basement membrane labeled with collagen-IV (red) in the indicated region of the flow conditioned sample in (C). Arrowheads indicate merging vessel walls in both two-dimensional and cross-sectional images. Vessel masks and fluorescent images represent at least three independent samples. Data represent mean ± S.E.M. Ordinary One-way ANOVA with Tukey's multiple comparisons test was performed; ns p > 0.05; **p < 0.01; ***p < 0.001; ****p < 0.0001 (n ≥ 3).

Our real-time observations of flow-conditioned and static vessels under high-VEGF conditions revealed an intriguing phenomenon of aberrant vessel growth occurring exclusively under flow. Under high-VEGF flow-conditioning, not only was flow-induced remodeling diminished, but the vessels also exhibited significant overgrowth (Figs. 6C, E and Supplementary Movie S2), resulting in an on average 28% greater increase in vessel coverage (from day 9 to day 12) compared to static samples under low-VEGF conditions (Supplementary Fig. S3). Characterization of the overgrowing vascular phenotype in flow-conditioned samples by immunostaining on day 12 confirmed the presence of merging tubular structures due to excessive growth and fusion of vascular tubes while maintaining three-dimensional hollow lumen structures (Fig. 6F). In contrast, static samples under high-VEGF conditions displayed much slower vessel growth (Figs. 6C, E and Supplementary Movie S2), significantly smaller than those under low-VEGF (Supplementary Fig. S3), as quantified by vessel coverage. These results suggest that static vessels continuously exposed to high levels of VEGF experience lagging gradual growth during the late maturation phase from day 9 to day 12. This lagging growth was also reflected in the connectivity index *C*, as vessels cultured in high-VEGF medium showed an average reduction in connectivity index *C* of approximately 10% on day 9 compared to those in low-VEGF conditions (where VEGF was reduced to the baseline level from day 6 to day 9 to induce vessel maturation, Fig. 6A). This led us to conclude that the sequential reduction of VEGF to the baseline level of 0.5 ng/ml, which characterizes our low-VEGF condition, is critical for sustained vessel development in microfluidic chips.

To further investigate the role of tumor-mimicking, high-VEGF levels for the observed overgrowth in flow-conditioned samples and slowed gradual growth in static samples, we tested the neutralizing effect of the anti-cancer drug bevacizumab under high VEGF conditions. Vessels were exposed to high levels of VEGF until day 9, and then bevacizumab was added to the high VEGF medium from day 9 to day 12 (Fig. 6B). Bevacizumab is a humanized anti-VEGF monoclonal antibody that neutralizes VEGF and blocks its signal transduction through VEGF receptors. It has been shown to inhibit VEGF-induced angiogenic activities in human endothelial cells without causing cytotoxicity [43]. The dosage of bevacizumab was determined based on the reported ratio to VEGF in human endothelial cells to ensure effective blockage of VEGF-induced angiogenic activities [43]. Interestingly, adding bevacizumab to the high-VEGF medium completely blocked the excessive vessel growth of flow-conditioned networks. Further, flow-induced remodeling is rescued, evidenced by the presence of pruned branches (Fig. 6D) and a reduction in vessel coverage (Fig. 6E). This suggests that elevated VEGF levels are the primary driver of excessive vessel growth under flow conditions. Also, in static samples, the addition of bevacizumab rescued the gradual vessel growth phenotype (Figs. 6D, E), bringing it closer to that of the low-VEGF static samples (Figs. 4A, D).

As a control for high-VEGF conditions, we added bevacizumab to low-VEGF medium from day 9 to day 12 in both flow- and static-conditioned samples (Fig. 7A). Bevacizumab had no effect on the gradual vessel growth in static samples or on the flow-induced remodeling in flow-conditioned samples (Fig. 7B). Static samples showed a shift toward larger tube diameters, while flow-conditioned samples showed a shift toward smaller diameters (Fig. 7C), consistent with low-VEGF conditions alone. Morphometric analysis showed no significant differences between low-VEGF and low-VEGF plus bevacizumab conditions, with vessel density increasing in static samples and decreasing in flow-conditioned samples due to network remodeling (Figs. 7D, E, F).

**Figure 7.**
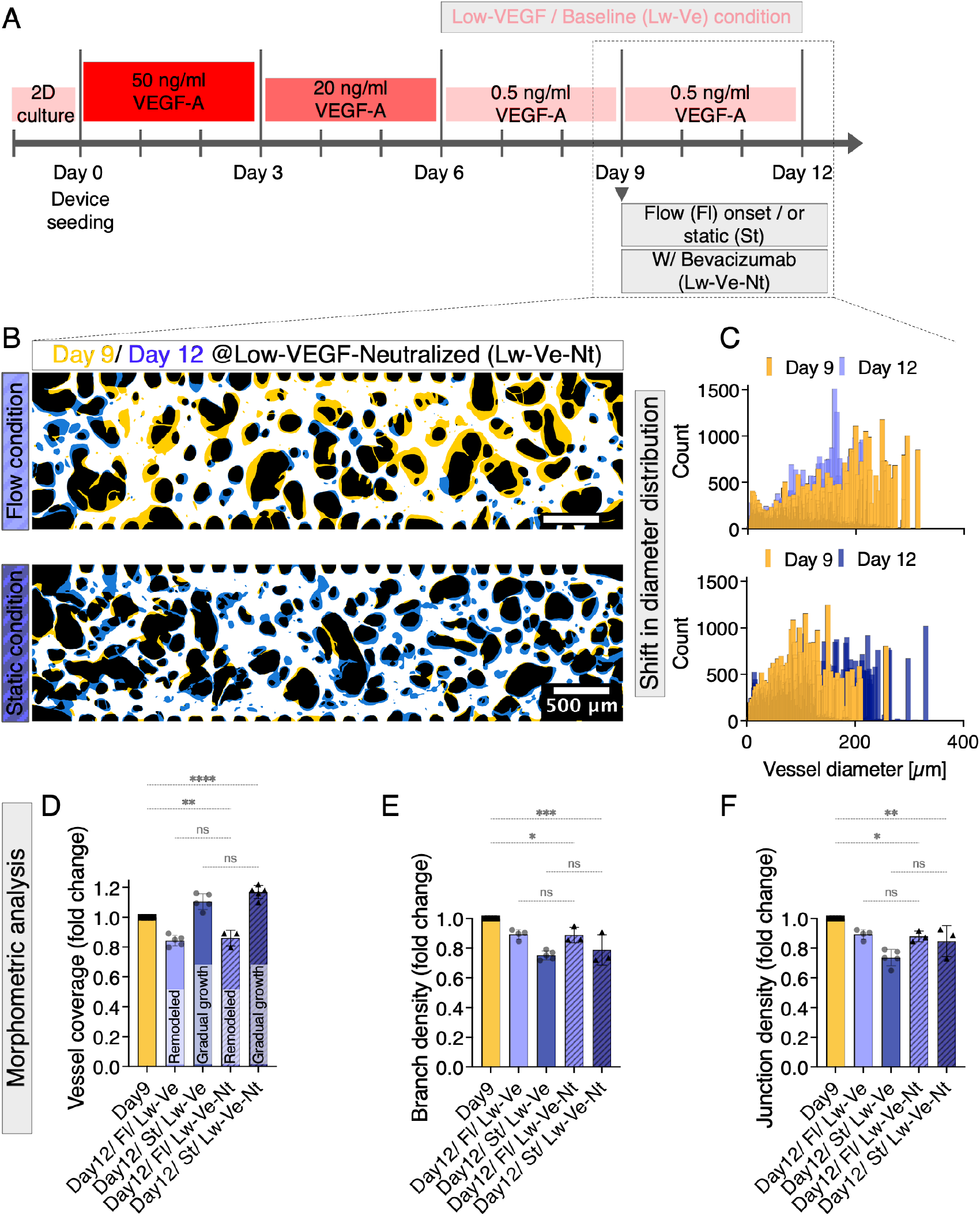
Cancer-drug induced VEGF neutralization has no impact on vascular remodeling under low-VEGF conditions. **(A)** Timeline of culture conditions from day 0 to day 12, with flow conditioning under low/baseline VEGF levels (Lw-Ve) with bevacizumab from day 9 to day 12. **(B)** Superimposed day 9 and day 12 masks showing pruning sites (yellow segments) under flow conditioning with low-VEGF plus bevacizumab as the perfusate, and enlarged vessels (blue segments) under static low-VEGF plus bevacizumab conditions. **(C)** Vessel diameter distribution on day 9 and day 12 for the flow conditioned and static samples shown in (B), which align with previous observations under low-VEGF conditions without bevacizumab (Fig. 4C). **(D-F)** Morphometric analysis underlines insignificant (p > 0.05) impact of bevacizumab under low-VEGF conditions (Lw-Ve-Nt) as quantified by **(D)** vessel coverage by area density, **(E)** vessel branching density as the number of branches per mm of vessel length, and **(F)** junction density as the number of junctions per mm of vessel length in day 12 vascular structures under low-VEGF (Lw-Ve, faded, same data points as in Figs. 4D, E, F) and low-VEGF plus bevacizumab (patterned, Lw-Ve-Nt) conditions for both flow conditioned (Fl) and static (St) samples, shown as fold change compared to day 9 values. Data represent mean ± S.E.M. Ordinary one-way ANOVA with Tukey's multiple comparisons test was performed; ns p > 0.05; *p < 0.05; **p < 0.01; ***p < 0.001; ****p < 0.0001 (n ≥ 3). Vessel masks are representative of at least three independent samples per group.

In conclusion, our results demonstrate that VEGF levels play a critical role in regulating flow-induced vascular remodeling. Under tumor-mimicking, high-VEGF conditions, flow-conditioned vessels exhibited significant overgrowth and blocked remodeling, suggesting excessive VEGF impairs proper vessel maturation. Static samples under high VEGF showed delayed growth, further highlighting the need for controlled VEGF reduction to sustain vascular development. VEGF neutralization with bevacizumab effectively blocked aberrant growth under flow conditions and restored remodeling in flow-conditioned networks. It also normalized vessel growth in static samples. Our findings highlight the importance of carefully adjusting the dose and duration of anti-VEGF therapies based on tumor vascularity and local VEGF fluctuations.

## Discussion

The interplay between VEGF signaling and blood flow is critical for vascular remodeling. This is particularly evident in pathological conditions such as tumor vascularization, which is characterized by tortuous, leaky, and dysfunctional vessels [2]. These abnormalities are driven by the excessive release of VEGFs by tumor cells, associated with defective vascular maturation and remodeling [1], [2], [18]. Combination therapies with anti-VEGF agents, such as bevacizumab, aim to normalize these vessels by restoring the balance between pro- and anti-VEGF signaling, thereby improving the delivery of chemo- and immunotherapeutic agents to the tumor [21]. However, clinical outcomes remain inconsistent [24], highlighting the need to better understand how flow dynamics and VEGF signaling drive vessel normalization. Using a vasculature-on-chip platform, we observed distinct effects of VEGF levels and flow on vascular remodeling. Under low-VEGF conditions, flow-induced remodeling refined network architecture and increased flow velocities. In contrast, static cultures without flow exhibited continuous overgrowth, resulting in a diminished network phenotype and reduced hypothetical flow velocities. Under tumor-mimicking, high-VEGF conditions, excessive overgrowth disrupted network refinement through remodeling. However, VEGF neutralization with the anti-cancer drug bevacizumab restored vessel remodeling. These findings underscore the intricate relationship between VEGF signaling and flow and provide valuable insights for improving anti-angiogenic therapies.

We took a quantitative approach to designing our vasculature-on-chip platform with optimized geometry and flow parameters to study vascular remodeling under controlled flow. COMSOL simulations ensured uniform flow distribution and long-term perfusability by adjusting the number of inter-post openings and the interface length between the media channels and the vascular chamber. The reduced chamber height (80 µm) confined the vessels to a planar morphology, preserving a hierarchy of vessel diameters (5-300 µm) despite the flattening of larger tubes. This planar setup enabled precise network quantification and computational flow simulations, which were critical for optimizing culture conditions and assessing the effects of VEGF and an anti-angiogenic drug. By isolating endothelial cells from stromal cells, we generated vascular tubes composed entirely of endothelial cells, simplifying initial observations of flow response and simulating conditions typical of aggressive tumors. Remarkably, these bare endothelial tubes exhibited flow-induced pruning, similar to remodeling observed in the mouse retinal vasculature [40], despite the absence of contractile wall-supporting cells such as pericytes. This aligns with previous reports under static conditions showing that on-chip vascular development is independent of the presence of pericytes [44]. Interestingly, while pericytes were found to significantly influence vessel diameter dynamics, a long-term reduction in vessel size was observed under static conditions regardless of their presence [44]. In contrast, our observations showed a long-term, unchecked gradual increase in vessel size under static conditions without pericytes, highlighting the need for future investigations into the role of pericytes under our modified condition of gradually reduced VEGF to better understand their impact on vascular remodeling.

To mimic physiological conditions, we identified reduced VEGF levels as essential for the transition of vessels from a rapid growth phase to an early maturation phase prior to flow initiation to assess flow-induced remodeling. For this early maturation phase (day 6 to day 9), we selected VEGF levels comparable to those found in healthy human blood and tissue [36], using 0.5 ng/mL as our baseline concentration. This approach allowed for near-plateau network growth by day 9, ensuring sufficient vessel connectivity for flow initiation. Flow-induced remodeling subsequently resulted in reduced vessel branching and more refined vessel structure. In contrast, a recent study initiated flow at day 6, resulting in increased vessel branching [44]. We speculate that vessels at day 6 remain in a phase of connection formation, while initiating flow at later stages, as in our approach, promotes enhanced network stabilization. Future studies could investigate how the timing of flow initiation affects the dynamics of vascular remodeling. In addition, further investigation of the interplay between flow mechanics and the growth and pruning patterns of our on-chip vasculature could provide deeper insights into how vascular remodeling shapes network hierarchy over time. Understanding how flow dynamics influence vascular organization and the efficiency of blood transport will advance knowledge of the mechanisms underlying vascular development and maturation. In addition to elucidating how remodeling transforms vessels from dense, disorganized plexuses to hierarchical, efficient structures, this understanding also enhances our ability to engineer advanced vascular networks that better meet the vascularization requirements of human organ-like and 3D tissue constructs.

Our gradual VEGF reduction strategy also highlights important differences from studies using commercial media with higher VEGF levels (4-5 ng/mL) [39], [44], [45]. These studies reported decreased vessel coverage under static conditions, whereas we observed gradual growth and increased coverage. Interestingly, our arrested vessel overgrowth under high VEGF conditions in static samples is consistent with findings of these studies suggesting that elevated VEGF levels compared to physiological levels in on-chip systems are critical. Differences in vessel coverage may also be due to media evaporation, which increases VEGF concentrations in their open-chip setups. To prevent this, we used an incubator microscope and housed the PDMS chips in chambered microscope slides with lids to minimize evaporation. This setup effectively minimized media evaporation and ensured consistent VEGF levels, which are critical for maintaining vessel maturation, connectivity, and flow response. Future studies could more precisely dissect the effects of a wider range of VEGF concentrations on vessel maturation and dysfunctional remodeling. Although we did not specifically test the 4-5 ng/mL range (our high-VEGF condition was 20 ng/mL), our results suggest that the dynamics of VEGF concentration over time significantly influence both flow-induced remodeling and gradual growth in static samples.

Our vascular chip provides a reliable platform to study tumor-like vascular phenotypes under high VEGF conditions characterized by excessive vessel overgrowth and disrupted network hierarchy. Bevacizumab treatment effectively reversed these abnormalities and restored flow-induced remodeling. These findings underscore the potential of the platform to test anti-angiogenic drugs and to study tumor vascular responses. Future studies could extend this approach to investigate a broader range of anti-angiogenic agents with different mechanisms of action, deepening our understanding of therapeutic outcomes and optimizing treatment strategies.

Our live imaging study of on-chip vasculogenesis highlights the complex interplay between VEGF signaling and flow dynamics in vascular remodeling, with important implications for anti-VEGF therapies. The results show that VEGF neutralization by the anti-VEGF drug bevacizumab effectively halts aberrant vascular growth and the resulting loss of network hierarchy under flow in high-VEGF conditions, thereby restoring flow-induced vascular remodeling. Consistent with vascular normalization theories for tumor vasculature [24], our results show that bevacizumab counteracts excessive tumor vascular overgrowth driven by high VEGF levels, promoting a more balanced and functional vascular network and potentially enhancing drug delivery to tumors. Conversely, under low-VEGF conditions, VEGF neutralization had no effect on the remodeling process, highlighting the context-dependent nature of anti-VEGF therapies. These results suggest that while anti-VEGF treatments are critical for controlling excessive angiogenesis, their effectiveness in stabilizing vascular networks depends on the local VEGF environment and tumor vascularity, particularly its perfusion levels. This underscores the need for personalized treatment strategies that take into account the specific VEGF context to optimize therapeutic outcomes. Future research could incorporate additional elements of the tumor microenvironment, such as immune cells and patient-derived cells, to further refine the model and enhance the exploration of personalized treatment strategies.

## Funding

This work was supported by the TUM International Graduate School of Science and Engineering (IGSSE), the Max Planck Society, funding from the European Research Council (ERC) under the European Union’s Horizon 2020 research and innovation program (Grant Agreement No. 947630, FlowMem) to K.A., and by the Federal Ministry of Education and Research (BMBF), the Free State of Bavaria under the Excellence Strategy of the Federal Government and the Länder for the TUM Innovation Network ''Robot Intelligence in the Synthesis of Life'' to A.R.B., F.C.S., K.A.

## Acknowledgments

The authors thank Dr. John Henningsen, Dr. Thomas Frank, and Dr. Louis Givelet for training in cleanroom techniques. We also thank Prof. Dr. Oliver Lieleg for access to shared cell culture facilities. We further thank Prof. Dr. Gabriel Amselem for his valuable insights on microfluidic perfusion techniques. This paper was typeset with the bioRxiv word template by @Chrelli: www.github.com/chrelli/bioRxiv-word-template

## Author contributions

Conceptualization, F.M.-S.; methodology, all authors, with significant contributions from F.M.-S. and K.A.; investigation, F.M.-S., E.H., and L.K., with significant contributions from F.M.-S.; resources, A.R.B., F.C.S., and K.A.; visualization: F.M.-S.; data curation: F.M.-S., J.T., and L.K.; writing original draft, F.M.-S. wrote the manuscript, and all authors commented on the manuscript and contributed to the writing process. supervision, F.M.-S. and K.A.; funding acquisition, A.R.B., F.C.S. and K.A.

## Competing interest statement

The authors declare no competing interests.

## Data and materials availability

The experimental data, chip design, and simulation data used in this study will be publicly available online at https://doi.org/10.14459/2024mp1762219

## Materials and Methods

### Experimental Design

The objective of this study was to develop a human vascular tissue-on-chip system optimized for studying the effects of long-term controlled flow on vascular network remodeling using real-time imaging. Additionally, the study investigated the impact of varying levels of VEGF (vascular endothelial growth factor) and the anti-VEGF antibody bevacizumab on vascular network formation and stability. The study's key components included fabricating microfluidic chips compatible with controlled flow conditioning via high-precision syringe pumps, culturing human endothelial and stromal cells within these chips, and applying controlled flow rates to simulate blood flow using perfusates at both baseline and elevated VEGF levels. Real-time imaging and quantitative analysis were employed to assess morphological and functional changes in the vascular networks in a three-day perfusion period. Predefined endpoints included evaluating network formation, stability, remodeling, and simulating flow distribution patterns in both empty chips and vascular networks before and after the perfusion experiment.

### Cell lines

Primary human umbilical vein endothelial cells (HUVECs, pooled donors, cryopreserved by PromoCell) and human pulmonary fibroblasts (HPFs, cryopreserved by PromoCell) were cultured in PromoCell endothelial growth medium-2 (EGM-2) and fibroblast growth medium-2 (FGM-2), respectively. In preparation for the chip seeding experiments, HUVECs and HPFs were cultured in culture dishes coated with collagen-I (Corning® BioCoat®) to approximately 80% confluency at passage 4 and passage 5, respectively, in a standard incubator at 37°C, 5% CO2 and 100% humidity.

### Microfluidic chip fabrication

Soft lithography techniques were used in class 4 clean room to produce negative silicon molds using two-inch silicon wafers (SIEGERT WAFER). SU-8 3000 photoresists (Microresist Technology GmbH) were used under appropriate spin-coating conditions to achieve a thickness of 80 µm. For casting, polydimethylsiloxane (PDMS, SYLGARD® 184 Elastomer Kit, Dow Corning) was used. The elastomer base and the elastomer curing agent were mixed 10:1. PDMS was degassed for 30 min and cured for 2 h at 80°C. After punching 1.5 mm inlet and outlet holes in PDMS slabs, air plasma (Zepto, Diener electronic GmbH & Co KG) was used for 60 s to facilitate the covalent bonding of PDMS to glass. The PDMS chips were fabricated on glass bottom µ-slides (ibidi GmbH), which were annealed at 70°C for 12 h to regain hydrophobicity after plasma treatment and sterilized using UV light for 1 h before cell seeding.

### Cell seeding and culture conditions

HUVECs and HPFs culture dishes were treated with cell detachment solution Accutase (Accutase®, Sigma-Aldrich) for 5-10 minutes and resuspended in phosphate-buffered saline (PBS) mixed with 1.6 U/ml thrombin (bovine plasma thrombin, Sigma-Aldrich) at concentrations of 20 million/ml and 8 million/ml, respectively. The cell suspension mixture was then mixed 1:1 with a fibrinogen mixture containing 8 mg/ml fibrinogen (bovine plasma fibrinogen, Sigma-Aldrich), 0.4 mg/ml collagen-I (collagen type I, rat tail, ibidi GmbH) and 0.4 U/ml aprotinin (MP Biomedicals™) and immediately injected into microfluidic chips for gelation. After injection, the chips were left on a flat surface at room temperature for 5 minutes and then placed in a CO2 incubator for 30 minutes before culture media was added. To mimic vascular maturation in culture, we sought an endothelial growth medium free of unknown components that could potentially prevent vascular maturation or cause experimental irreproducibility. The EGM-2 medium used in this study has the lowest level of bovine serum (0.02%) on the market and contains no other unknown ingredients. The manufacturer reports the exact levels of each ingredient, including VEGF-A, which is provided at 0.5 ng/mL, the lowest commercially available concentration to our knowledge. The amount of medium added to each chip followed the manufacturer's recommendations, taking into account the number of cells per chip. After network formation on day 6, the medium was replaced every 24 hours. We estimate the time scale of nutrient diffusion through the gel to be approximately 1.6 hours to reach a homogeneous concentration throughout the vascular chamber. This is based on the diffusivity of a protein of comparable size to VEGF-A (approximately 40 kDa) in a 4 mg/ml fibrin gel, which is close to its diffusivity in water (approximately 10^-6 cm^2^/s) [46], [47], and the geometry of our device, i.e., the half-width of the vascular chamber through which molecules must diffuse into the fibrin gel to feed the cells. To estimate the diffusion time of molecules in our chips, we used a simplified approach based on Fick's second law of diffusion, where the diffusion time is determined by dividing the square of the diffusion distance by the diffusivity.

In our optimized vascular maturation mimicking culture protocol, EGM-2 supplemented with 0.2 U/ml aprotinin and 50 ng/ml VEGF-A (human VEGF165, ReliaTech), referred to as the “pro-angiogenic VEGF-A level,” was initially introduced into the media channels and reservoirs. On day 3, the media was replaced with EGM-2 + 0.2 U/ml aprotinin + 20 ng/ml VEGF-A (“high VEGF-A level”) and then switched to EGM-2 + 0.2 U/ml aprotinin (“low/baseline VEGF-A level”) on day 6. This baseline medium was refreshed every 24 hours until day 9. In our tumor vasculature mimicking culture protocol, the vascular chips were similarly treated with pro-angiogenic medium until day 3, followed by high-VEGF medium from day 3 to day 6. However, instead of switching to the baseline medium, the high VEGF-A medium was maintained from day 6 to day 9, with media refreshed every 24 hours.

### Controlled flow and real-time imaging

For flow experiments, vascular chips were flow-conditioned from day 9 to day 12. Prior to perfusion, networks were assessed for perfusability by FITC-Dextran (500 kDa, Sigma-Aldrich) infusion on day 8. Visualization of the dextran infusion was combined with quantification of the connectivity index *C* to confirm the perfusability of the vascular networks (Figs. 1C, D). Under optimized vascular maturation mimicking culture conditions, an average connectivity index *C* of 0.86 ensured vessel stability throughout the perfusion experiments. In contrast, under tumor vessel mimicking conditions, an average connectivity index *C* of 0.79 was sufficient to maintain vessel stability throughout the experiments.

The perfusion circuit was set up on day 8 with the fluids kept inside the CO2 incubator and the pump modules outside. All perfusate mixtures were prepared on day 8 and kept in the CO2 incubator for 24 h before the perfusion experiment. The CETONI Nemesys S low-pressure pump system was used for perfusion, which allows precise sub-nanoliter fluid dosing. Low-pressure 10 ml glass syringes (SETonic GmbH, Germany) were mounted on a pump module (CETONI GmbH, Germany) per microfluidic chip controlled by CETONI Elements software. 1.5 mm Teflon tubing (TECHLAB GmbH) and flangeless fittings (IDEX Health & Science LLC) connected the chips to the pump modules. Homemade PDMS pins were used to block the injection holes. Teflon tubing was inserted directly into the chip inlet and outlet holes. Like the perfusate, the medium for the static samples was maintained in the CO2 incubator throughout the perfusion experiment and was replaced every 24 hours with a volume equal to the total amount of medium perfused through the flow-conditioned samples during the same period. To evaluate the effects of VEGF levels under static and flow conditions, both high- and low-VEGF and high- and low-VEGF plus bevacizumab perfusates were prepared and incubated for 24 hours prior to the perfusion experiment. Bevacizumab was added to the high-VEGF medium at a molar ratio of 2.6:1 bevacizumab to VEGF-A.

Live brightfield imaging in the CO2 incubator was performed with Etaluma LS720 to monitor vessels over time. A µ-Slide microscopy rack with magnetic lids (ibidi GmbH) was used to stabilize flow- and static-conditioned samples and to prevent evaporation of medium in static samples over the 3-day course of live imaging.

### Immunofluorescence staining

For immunostaining, samples were fixed with 4% paraformaldehyde (PFA) in PBS for 15 minutes at room temperature on day 12. After fixation, the samples were permeabilized with 0.1% Triton X-100 in PBS for 15 minutes at room temperature. After permeabilization, the samples were incubated overnight at 4°C with primary antibodies anti-collagen IV (eBioscience™, 51-9871-82, 4µg/ml) and anti-ICAM-2 (eBioscience™,11-1029-42, 2µg/ml) in PBS with 0.01% bovine serum albumin (BSA). After primary antibody incubation, the samples were washed with PBS and then incubated with 0.1 µg/ml DAPI in PBS for 10 minutes at room temperature to stain the nuclei. Finally, the chips were washed and immersed in PBS for maintenance and imaging. Stained vasculatures were visualized using a confocal laser scanning microscope (Leica SP8).

### Image processing and morphometric analysis

Brightfield and fluorescent images were pre-processed using the open-source platform for biological image analysis Fiji (ImageJ2, version 2.14.0/1.54f) [48]. The avascular gaps between the vessels were first manually segmented in the brightfield images as “negative masks” to be trained into a deep learning-based model with Cellpose [49], [50]. Then the vessel masks were extracted by taking the inverse of the negative masks. Key metrics such as vessel coverage, branching density, and junction density were calculated based on these segmented vessel masks. Morphometric analysis of the binarized vessel masks was performed using Fiji's Vessel Analysis [51], Skeletonize (2D/3D), and AnalyzeSkeleton (2D/3D) [52] plug-ins. Prism 10 GraphPad software was used for data visualization and plotting. To determine the connectivity index *C*, binarized vessel masks were processed in MATLAB for skeletonization and segmented into individual branches. Branches connected to the media channel through inter-post openings at the upper or lower interfaces of the vascular chamber were identified and annotated as connected nodes. For each node on the upper boundary, the shortest path length *Li* to any node on the lower boundary was computed, and vice versa for lower nodes to upper nodes. Each shortest path *Li* was then normalized to the channel width W to calculate the ratio *W/ Li*. The connectivity index *C* was obtained by summing these ratios and dividing by the total number of potential openings (48 per chip: 24 opposite pairs, Fig.1D).

### Computational fluid dynamics simulation

To analyze the fluid dynamics within the vascular networks, two-dimensional vessel masks obtained from image segmentation were converted to drawings using Inkscape software. These DXF drawings were then pre-processed and scaled using AutoCAD to ensure accurate dimensions and compatibility with subsequent computational modeling. The processed vessel geometries were imported into a computational fluid dynamics solver (COMSOL Multiphysics, version 6.0). The flow field within the vascular networks was determined by solving the Navier–Stokes equations under the assumptions of two-dimensional laminar and incompressible flow with no-slip boundary conditions using a finite element method in COMSOL. The inlet boundary condition was defined with an inflow velocity of 80.13 µm/s, corresponding to the experimental volumetric flow rate of 0.5 µl/min, an inlet width of 1.3 mm, and a chip height of 80 µm. The outlet boundary condition was configured to ensure flow continuity by matching the inflow. Literature values for cell culture medium density (998.2 kg/m3) and dynamic viscosity (0.94 mPa.s) were used to accurately represent the physical properties of the fluid [39]. Simulated velocity fields and velocity distribution plots were derived from COMSOL simulations. Velocity field data were exported from the vessel region between the upper and lower openings on a uniform grid matched to the resolution of the imaging data. The simulated velocity field was normalized using the Darcy velocity calculated for each network in Matlab. In our vascular networks, the Darcy velocity is determined by dividing the inflow rate of 0.5 µl/min by the product of the cross-sectional area and the porosity. Porosity is calculated as the ratio of vessel area to total area. Matlab was used to plot velocity distributions.

### Statistical Analysis

All statistical analyses were performed using Prism 10 GraphPad software. Appropriate statistical significance tests were selected based on the experimental design and data distribution. For experiments comparing multiple groups, ordinary one-way ANOVA was performed, followed by Tukey's multiple comparison test. Data are presented as mean ± S.E.M., with 95% confidence intervals. At least three independent biological replicates (n ≥3) were used for each experiment. All statistical analyses for experimental data were performed on at least three independent samples per condition.

## Supplementary Materials

**Supplementary Figure S1.**
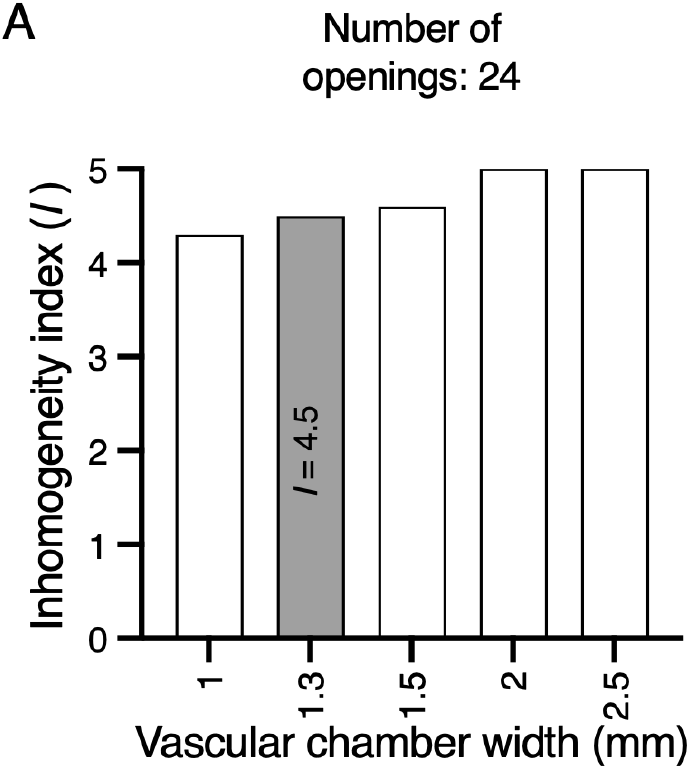
Effect of vascular chamber width on inhomogeneity index. **(A)** Inhomogeneity indices I calculated from simulated velocities in chips with different vascular chamber widths, where the number of openings was fixed at 24 and the media channel width was kept constant at 1.3 mm.

**Supplementary Figure S2.**
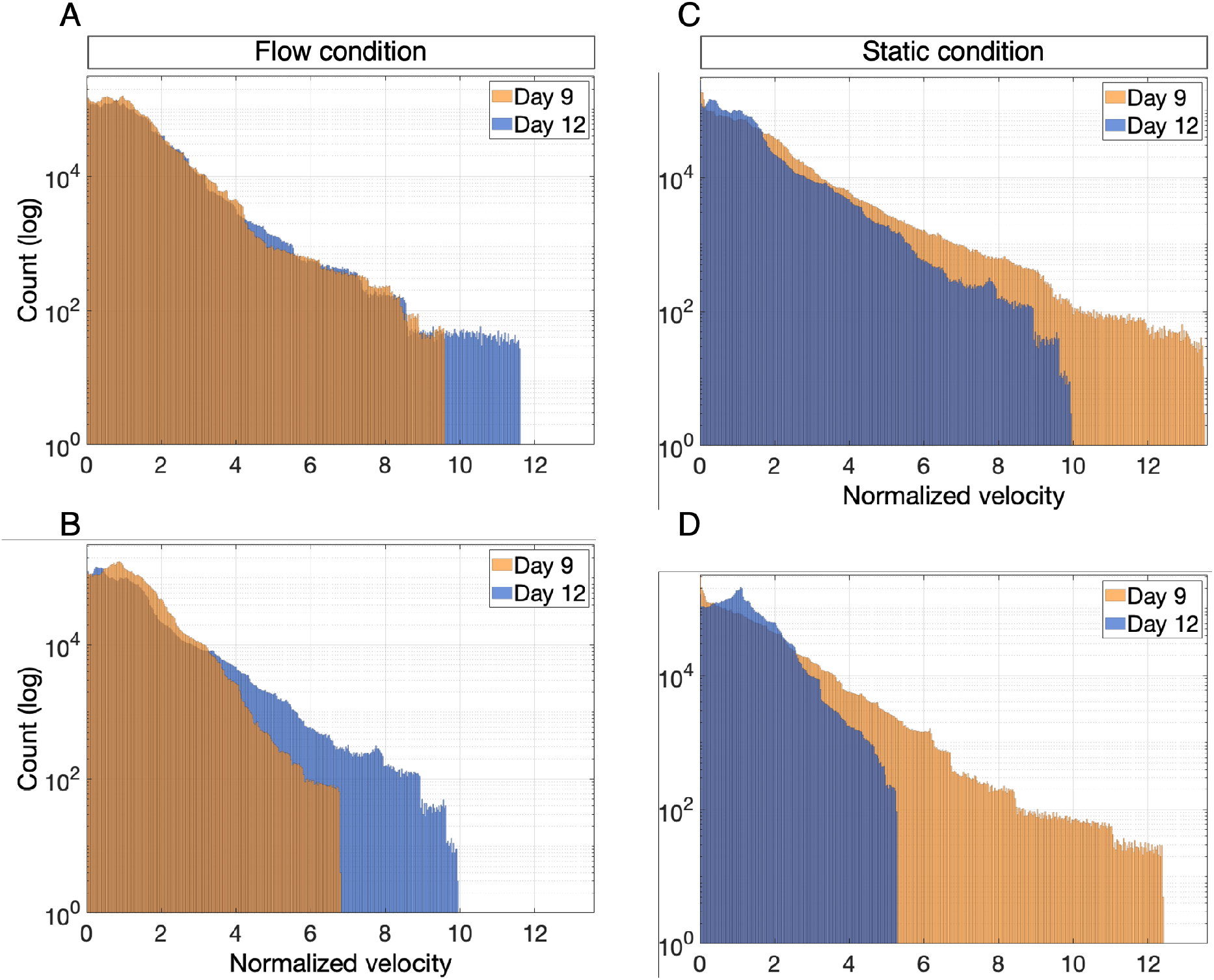
Flow-induced network remodeling consistently shifts normalized velocity distributions toward higher velocities across multiple independent samples. **(A-D)** Velocity distribution plots for the second and third independent static and flow-conditioned samples show trends consistent with Figs. 5B and 5C. **(A, B)** Normalized velocity distribution (by Darcy velocity) for the second and third independent flow-conditioned samples (Chip2 and Chip3 in Fig. 5D) show a shift toward higher velocities on day 12 compared to day 9. **(C, D)** Normalized velocity distribution (by Darcy velocity) for the second and third independent static samples (Chip5 and Chip6 in Fig. 5D) show a shift toward lower velocities on day 12 compared to day 9. Note that the visualized velocity fields for day 9 and day 12 in the static samples were artificially modeled because these samples were not exposed to flow conditions.

**Supplementary Figure S3.**
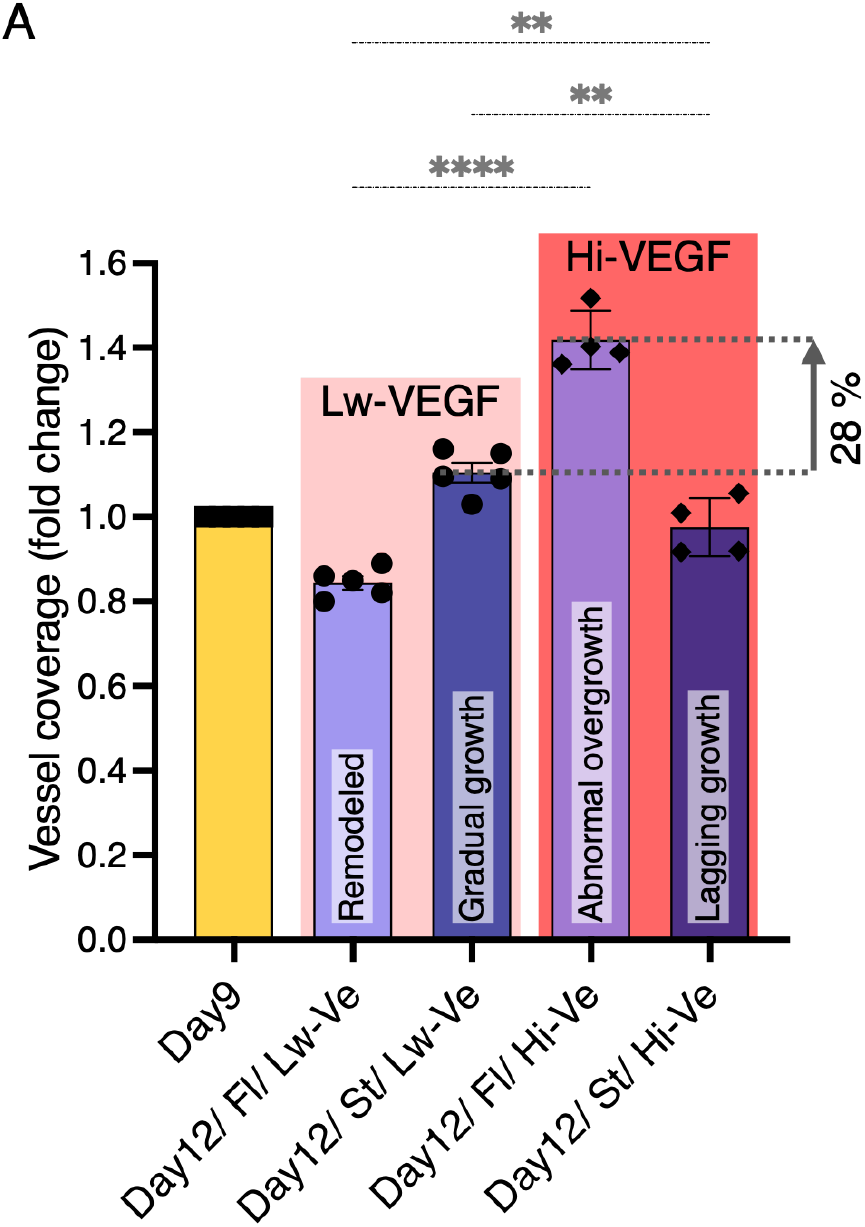
Effect of VEGF levels on vessel coverage. **(A)** Quantification of vessel coverage by area density under low-VEGF (Lw-Ve, same data points as in Fig. 4D) and high-VEGF (Hi-Ve, same data points as in Fig. 6E) conditions for both static (St) and flow-conditioned (Fl) samples at day 12, presented as fold change relative to day 9 values. Group comparison shows that, compared to low-VEGF, high-VEGF caused a significant abnormal overgrowth in flow-conditioned samples and a significantly lagging vessel growth in static samples. This significant overgrowth underflow conditions resulted in an average of 28% greater increase in vessel coverage compared to the gradually growing static samples cultured under low-VEGF conditions. Data are shown as mean ± S.E.M. Ordinary one-way ANOVA followed by Tukey's multiple comparison test was performed; **p < 0.01; ****p < 0.0001 (n ≥ 4).

**Supplementary Movie S1. Low-VEGF: vessel remodeling under flow conditions and gradual vessel growth under static conditions**.

This time-lapse movie captures the evolution of the same vascular networks as shown in Fig. 4A underflow and static conditions, with images taken at 6-hour intervals. Under flow conditions, the perfused network responds dynamically to flow, with vessels undergoing wall rearrangement and selective branch pruning. These flow-induced changes reduce vessel coverage (Fig. 4D) and slightly decrease branching and junction density (Figs. 4E, F), demonstrating the remodeling process driven by continuous flow. In contrast, under static conditions, gradual vessel growth leads to increased vessel coverage (Fig. 4D), which results in a significantly larger decrease in branching and junction density (Figs. 4E, F) due to lateral fusion of the vascular tubes, highlighting the instability and reduced hierarchy of the network without flow. The movies combine bright-field images with segmented vessel masks, where avascular gaps in the network appear as white-shaded background regions. Movies represent at least three independent samples.

**Supplementary Movie S2. High-VEGF: aberrant vessel overgrowth under flow conditions and lagging gradual growth under static conditions**.

This time-lapse movie captures the behavior of the same vascular networks as shown in Fig. 6C under high-VEGF conditions in both flow- and static-conditioned samples, with images taken at 6-hour intervals. In flow-conditioned samples, high-VEGF exposure leads to aberrant vessel overgrowth and reduced flow-induced remodeling. This overgrowth results in a 28% greater increase in average vessel coverage than static samples under low-VEGF conditions (Fig. S3), highlighting the excessive expansion caused by high VEGF during perfusion. In contrast, high-VEGF static samples show delayed, gradual vessel growth, significantly slower than low-VEGF static samples (Fig. S3). This suggests that vessels exposed to high VEGF experience delayed growth during the late maturation phase, from day 9 to day 12. The movies combine bright-field images with segmented vessel masks, where avascular gaps in the network appear as white-shaded background regions. Movies represent at least three independent samples.

## Notes

https://doi.org/10.14459/2024mp1762219

### Competing Interest Statement

The authors have declared no competing interest.

### Summary of Updates

abstract shortened, results section and Figure 2 updated

